# Global Progress in Competitive Co-Evolution: a Systematic Comparison of Alternative Methods

**DOI:** 10.1101/2024.06.06.597852

**Authors:** Stefano Nolfi, Paolo Pagliuca

## Abstract

We investigate the use of competitive co-evolution for synthesizing progressively better solutions. Specifically, we introduce a set of methods to measure historical and global progress. We discuss the factors that facilitate genuine progress. Finally, we compare the efficacy of four qualitatively different algorithms. The selected algorithms promote genuine progress by creating an archive of opponents used to evaluate evolving individuals, generating archives that include high-performing and well-differentiated opponents, identifying and discarding variations that lead to local progress only (i.e. progress against a subset of possible opponents and retrogressing against a larger set). The results obtained in a predator-prey scenario, commonly used to study competitive evolution, demonstrate that all the considered methods lead to global progress in the long term. However, the rate of progress and the ratio of progress versus retrogressions vary significantly among algorithms.

## 1. Introduction

Recent advances in machine learning have demonstrated the importance of using large corpora of training data. In the context of embodied and situated agents, this implies placing the agents in complex and diversified environments. However, manually designing environments of this kind is difficult and expensive. A convenient alternative is constituted by multi-agent scenarios in which adaptive agents are located in environments containing other adaptive agents with conflicting goals—a method referred to as competitive co-evolution (Simione & Nolfi, 2021) or self-play (Bansal et al., 2017). In these settings, the learning agents are exposed to environmental conditions that vary continuously as a result of the behavioral modifications of the other adaptive agents. In other words, these settings permit the automatic generation of a large corpus of training data.

Competitive settings also offer other important advantages. They can spontaneously produce convenient learning curricula (Wang et al., 2021) in which the complexity of the training conditions progressively increases while the skills of the agents—and consequently their ability to master complex conditions—also improve. Finally, such settings may naturally generate a form of adversarial learning (Biggio & Roli, 2018), where the training data are shaped to challenge the weaknesses of the adaptive agents.

Unfortunately, competitive co-evolution does not necessarily result in a progressive complexification of agents’ skills or environmental conditions. As highlighted by Dawkins and Krebs in the context of natural co-evolution (Dawkins & Krebs, 1979), the evolutionary process can lead to four distinct long-term dynamics: (1) extinction: one side may drive the other to extinction, (2) definable optimum: one side might reach a definable optimum, preventing the other side from reaching its own optimum, (3) mutual local optimum: both sides may reach a mutual local optimum, and (4) endless limit cycle: the race may persist in a theoretically endless limit cycle, in which similar strategies are abandoned and rediscovered over and over again.

Regrettably, pioneering attempts to evolve competing robots consistently yield the last undesirable outcome described by Dawkins and Krebs. Initially, there is true progress, but subsequently agents modify their strategies, resulting in apparent progress. In other words, they improve against the strategies exhibited by their current opponents while retrogressing against other strategies that are later adopted by opponents (Miconi, 2008). Consequently, limit cycle dynamics emerge, where the same strategies are abandoned and rediscovered repeatedly (Sinervo & Lively, 1996; Miconi, 2009; Nolfi, 2012).

The generation of genuine progress requires the usage of special algorithms that: (i) expose the adapting agents to both current and ancient opponents (Rosin & Belew, 1997), (ii) expose the adaptive agents to a diverse range of opponents (De Jong, 2005; Simione & Nolfi, 2021), and (iii) identify and retain variations that lead to true progress only (Simione & Nolfi, 2021). Furthermore, analyzing competitive settings necessitates the formulation of suitable measures to differentiate between true and apparent progress and to evaluate the efficacy of the obtained solutions.

In this article, we present a systematic comparison of alternative competitive co-evolutionary algorithms, including novel variations of state-of-the-art algorithms. Additionally, we describe the measures that can be used to analyze the co-evolutionary process, discriminate between apparent and true progress, and compare the efficacy of alternative algorithms.

## 2. Measuring progress

In evolutionary experiments in which the evolving individuals are situated in solitary environments, the fitness measured during individuals’ evaluation provides a direct and absolute measure of performance. Usually the initial state of the robot and of the environment is subjected to random variations, which impact on the fitness measure. However, these variations do not have an adversarial nature. Consequently, the impact these variations have can be usually reduced by evolving solutions that are robust with respect to them and by averaging the fitness obtained during multiple evaluation episodes (Carvalho & Nolfi, 2023, Pagliuca & Nolfi, 2019).

In competitive social settings, instead, the fitness obtained during individuals’ evaluation crucially depends on the opponent/s situated in the same environment. Consequently the method used to select opponents has a pivotal effect on the course of the co-evolutionary process.

The dependency of the fitness measure on the characteristics of the opponents also affects: (1) the identification of the best solution generated during an evolutionary process, (2) the estimation of the overall effectiveness of a solution, and (3) the comparison of the efficacy of alternative experimental conditions. The former two problems can be tackled by identifying a specific representative set of opponents known as “champions”, which typically include the best opponents produced during independent evolutionary experiments.

The third issue can be resolved through a technique called “cross-test”. In a cross-test, the best solutions obtained from N independent evolutionary experiments conducted under one experimental condition are evaluated against the best opponents obtained from N different independent evolutionary experiments.

Lastly, the dependence of the fitness measure on opponent characteristics impacts the method used to measure evolutionary progress. In non-competitive settings, progress and retrogression can be straightforwardly measured by computing the variation of fitness across generations. Instead, in competitive settings, measuring progress becomes more challenging.

As highlighted by Miconi (2009), we need to differentiate between three types of progress: (i) local progress, i.e. progress against current opponents, (ii) historical progress, i.e. progress against opponents of previous generations, and (iii) global progress, i.e. progress against all possible opponents. To measure local progress, we can post-evaluate agents against opponents from recent preceding generations. Historical progress can be assessed by evaluating agents against opponents from previous generations. These data can be effectively visualized using the “Current Individual against Ancestral Opponents” (CIAO) plots introduced by Cliff & Miller (1995, 2006). For global progress estimation, we can post-evaluate agents against opponents generated in independent evolutionary experiments—opponents that differ from those encountered during the evolutionary process. Additionally, an indication of global progress can be obtained by evaluating agents against opponents from future generations. The data obtained by post-evaluating agents against opponents of previous and future generations can be conveniently visualized using the master tournament plots introduced by Nolfi & Floreano (1998).

## 3. Competitive evolutionary algorithms

In this section we will review the most interesting co-evolutionary algorithms described in the literature and the methods that we will compare in our experiments.

We focus our analysis on evolutionary algorithms attempting to maximize the expected utility, i.e. the expected fitness against a randomly selected opponent or the average fitness against all possible opponents. Other researchers have explored the use of competitive evolution for synthesizing Nash equilibrium solutions (Figici & Pollack, 2003; Wiegand et al., 2002) and Pareto-optimal solutions (De Jong, 2004).

As previously mentioned, achieving true progress necessitates the utilization of specialized algorithms that: (i) expose adapting agents to both current and ancient opponents, (ii) evaluate adaptive agents against a diverse set of opponents, or (iii) identify and retain variations that lead to genuine progress.

The first method we consider is the **Archive** algorithm, as introduced by Rosin and Belew (1997), see also (Stolfi et al., 2021). In this algorithm, a copy of the best individual from each generation is stored in a “hall-of-fame” archive. Opponents are then randomly selected from this archive using a uniform distribution. Evaluating evolving agents against opponents from previous generations clearly promotes historical progress. While the production of global progress is not guaranteed, it can be expected as a form of generalization. Indeed, the need to defeat an increasing number of ancient opponents should encourage the development of strategies that generalize to opponents not yet encountered.

The second method that we will consider is the **Maxsolve\*** algorithm, a variation of the original method introduced by De Jong (2005), see also (Samothrakis et al., 2013; Liskowski & Krawiec, 2016). The Maxsolve algorithm operates by using an archive containing a predetermined maximum number of opponents. The size of the archive is kept bounded by removing dominated and redundant opponents from it. The domination criterion works for transitive problems, i.e. in problems where agents A outperforming agents B, which in turn outperform agents C, necessarily outperform agents C. To allow the algorithm to operate with non-transitive problems, like the predator and prey task considered in this paper, we implemented the Maxsolve* algorithm, a variation of the original Maxsolve algorithm, which retains in the archive the individuals achieving the highest performance, on average, against the opponents of the 10 preceding phases. The method thus attempts to automatically select high-quality champions that can promote the discovery of high-quality solutions and minimize the time spent evaluating agents against poor opponents (for a related approach see, Bari et al., 2018). Clearly, the size of the archive (AS) plays an important role in this method. Indeed, the smaller the size of the archive is, the smaller the training data is. On the other hand, the smaller the size of the archive is, the higher the minimization of the time spent against poor opponents is. By systematically varying the size of the archive (data not shown), we observed that the best results are obtained by using relatively large archives, i.e. archives containing up to 25,000 opponents.

The third method is the **Archive\*** algorithm, a novel approach introduced in this paper as a variation of the original Archive algorithm described earlier. The Archive* algorithm operates by evolving N independent populations, each contributing to the generation of a single archive. These evolving populations are then evaluated against opponents selected from the shared archive.

The rationale behind this method lies in the use of multiple populations, which enhances the diversification of opponents included in the archive. Specifically, the Archive* method automatically generates multiple families of diversified champions, representing alternative challenges for the evolving agent. The Archive* algorithm shares similarities with the population-based reinforcement learning method proposed by Jaderberg et al. (2018).

Finally, the fourth method we propose is the **Generalist** algorithm, introduced by Simione and Nolfi (2021). In this approach, agents are evaluated against a subset of opponents, while the remaining opponents serve to discriminate between agents who retained variations leading to global progress and those who retained variations leading to local progress. This information guides the preservation or discarding of individuals. The subset of opponents used for agent evaluation is randomly selected at regular intervals (every N generations) to maximize the functional diversity of the subset. Unlike the previous algorithms, the Generalist method does not rely on archives.

Another potential approach involves using randomly generated opponents (Chong et al., 2009, 2012; Jaśkowski et al., 2013). The primary advantage of this technique lies in the direct promotion of global progress because agents are consistently evaluated against new opponents. However, there is a significant drawback: the efficacy of these opponents does not improve over generations. Consequently, this method does not allow for the development of agents capable of defeating strong opponents. In fact, the performance obtained using this approach by Samothrakis et al. (2013) were considerably lower than the performance achieved with a variation of the Maxsolve algorithm described earlier.

The methods described above are meta-algorithms that should be combined with an evolutionary algorithm to determine how populations of individuals vary across generations. In previous studies, standard evolutionary algorithms or evolutionary strategies were employed. Instead, in this work, we utilize the OpenAI-ES algorithm (Salimans et al., 2017), which represents a “modern” evolutionary strategy (Pagliuca et al., 2020). The OpenAI-ES algorithm leverages the matrix of variations introduced within the population and the fitness values obtained by corresponding individuals to estimate the gradient of fitness. It then guides the population’s movement in the direction of this gradient using a stochastic optimizer (Kingma & Ba, 2014). Importantly, this algorithm is well-suited for non-stationary environments. This is because the momentum vectors also guide the population in the direction of previously estimated gradients, thereby enhancing the possibility of generating solutions effective against opponents encountered in previous generations.

For a detailed description of the algorithms, please refer to the Appendix A. Additionally, the experiments can be replicated using the evorobotpy2 tool available from the https://github.com/snolfi/evorobotpy2GitHub repository.

## 4. The predator and prey problem

We chose to compare alternative algorithms using a predator and prey problem because it represents a challenging scenario (Miller & Cliff, 1994). Additionally, this problem is widely used for studying competitive evolutionary algorithms (Miller & Cliff, 1994; Floreano & Nolfi, 1997a; Floreano & Nolfi, 1997b; Floreano et al., 1998; Nolfi & Floreano, 1998; Stanley & Miikkulainen, 2002; Buason & Zimke, 2003; Buason et al., 2005; Jain et al., 2012; Palmer & Chou, 2012; Ito et al., 2013; Lan et al., 2019; Georgiev et al., 2019; Lee et al., 2021; Simione & Nolfi, 2021; Stolfi et al., 2021).

The predator and prey problem presents highly dynamic, largely unpredictable, and hostile environmental conditions. Consequently, it necessitates the development of solutions that are fast, robust, and flexible. Moreover, agents must exhibit a range of integrated behavioral and cognitive capabilities, including avoiding fixed and moving obstacles, optimizing motion trajectories considering multiple constraints, integrating sensory information over time, anticipating opponent behavior, disorienting opponents, and adapting behavior on the fly based on the opponent’s actions (Humphries & Driver, 1970).

The robots in our study are simulated MarXbots (Bonani et al., 2010) equipped with neural network controllers. The connection strengths within the robots’ neural networks, which determine their behavior, are encoded in artificial genotypes and evolved. Specifically, predators are evolved to enhance their ability to capture prey (i.e., reach and physically touch the prey) as quickly as possible, while preys are evolved to maximize their ability to avoid being captured for as long as possible. The fitness of predators corresponds to the fraction of time required to capture the prey. The fitness of preys is the inverse of the fraction of time required by the predator to capture them.

A detailed description of the robots’ characteristics and their neural controllers can be found in the Appendix B.

To compare the relative effectiveness of the considered methods, we continued the evolutionary process until a total of 75 billion evaluation steps were performed.

For those interested in replicating the experiments, the evorobotpy2 tool is available at the https://github.com/snolfi/evorobotpy2GitHub repository.

## 5. Results

In this section, we present the results obtained using the Archive, Maxsolve*, Archive*, and Generalist algorithms. The results include data collected from 10 replication experiments conducted with each algorithm, resulting in a total of 40 experiments.

Figure 1 displays master tournament data, i.e. the performance of predator and prey robots of different generations evaluated against opponents of different generations. The results demonstrate that all considered methods exhibit historical progress overall. Notably, the robots perform better against opponents from previous generations than against those from successive generations in the majority of cases. Additionally, these data provide an indication of global progress, as the robots of generation X+N perform better against opponents from future generations than robots of generation X in most of the cases. However, it is worth noting that only the Generalist algorithm consistently produces progress across all phases, resulting in monotonically better robots. The other algorithms also exhibit retrogressions, albeit less frequently than progress.

**Figure 1.**
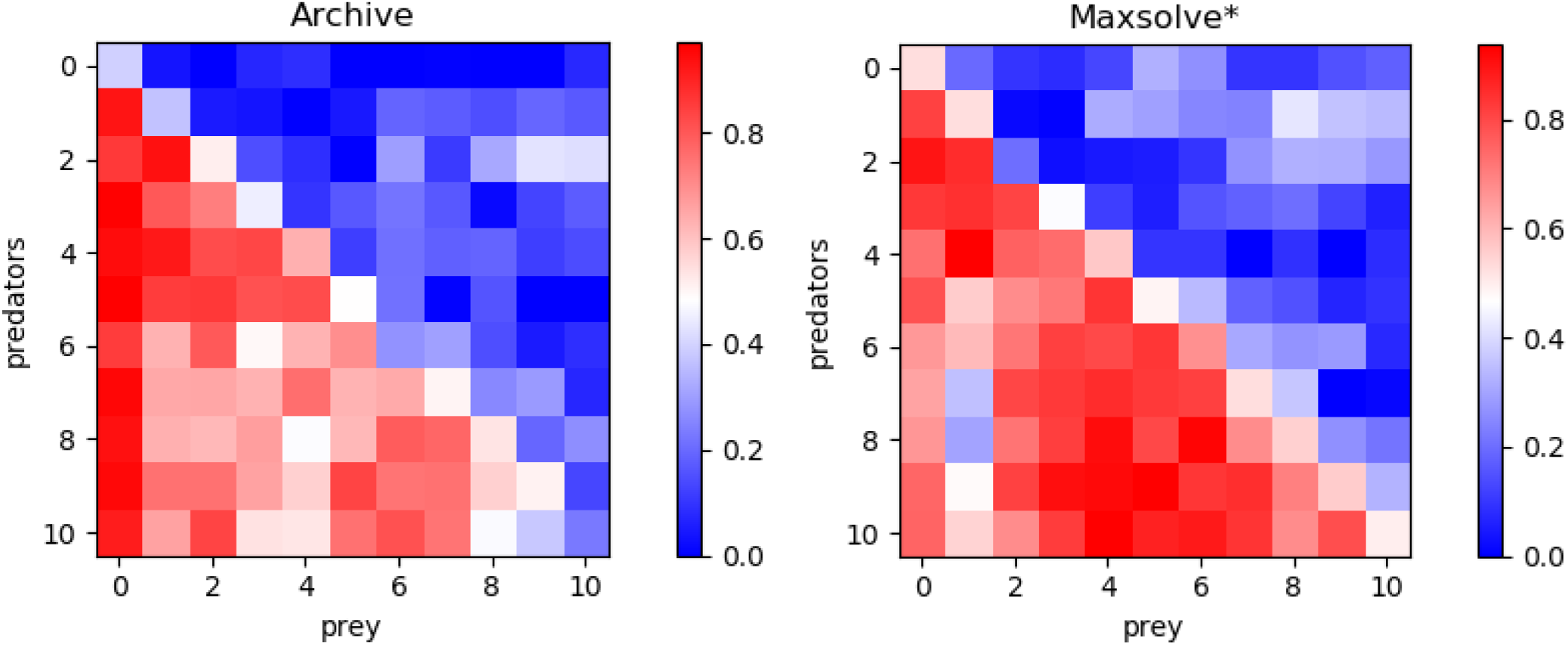
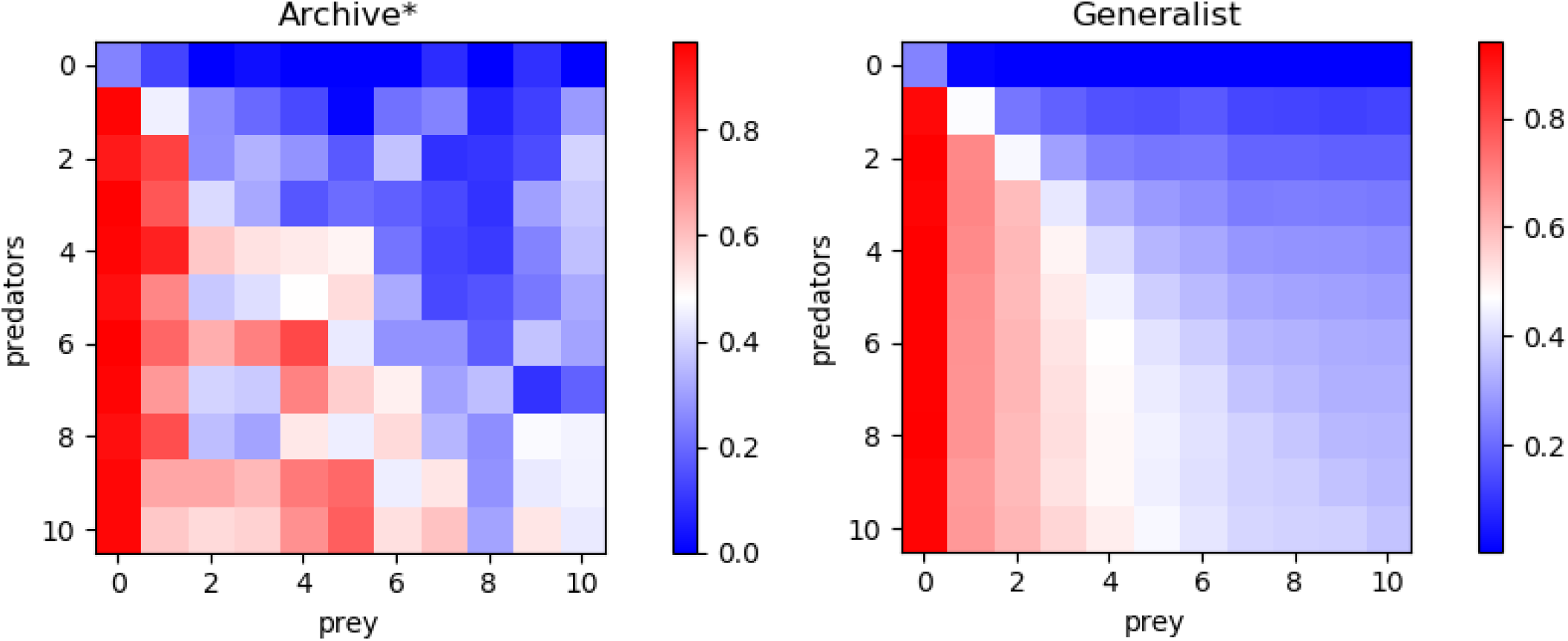
Performance of predators at different evolutionary stages, post-evaluated against opponents of all stages. Results obtained with the Archive, Maxsolve*, Archive* and Generalist algorithms. Data was collected by evaluating the predators of phase 0 (generation 0) and the subsequent 10 stages against prey of generation 0 and the following 10 stages. The stages are separated by 1/10 of the total generations. The performance of the prey corresponds to 1.0 minus predator performance. Each plot represents the average results of 10 evolutionary experiments.

This qualitative difference is confirmed by Figure 2, which illustrates the performance of the robots from last generation against opponents from both the current and previous generations. The Archive and Maxsolve* robots of the last generation consistently exhibit better and better performance against opponents of the last four preceding stages only. The Archive* robots of the last generations consistently display better and better performance against opponents of the last 3 stages (for predator robots) and the last 5 stages (for prey robots) only. Only robots evolved with the Generalist algorithm consistently exhibit improved performance against older opponents across all phases.

**Figure 2.**
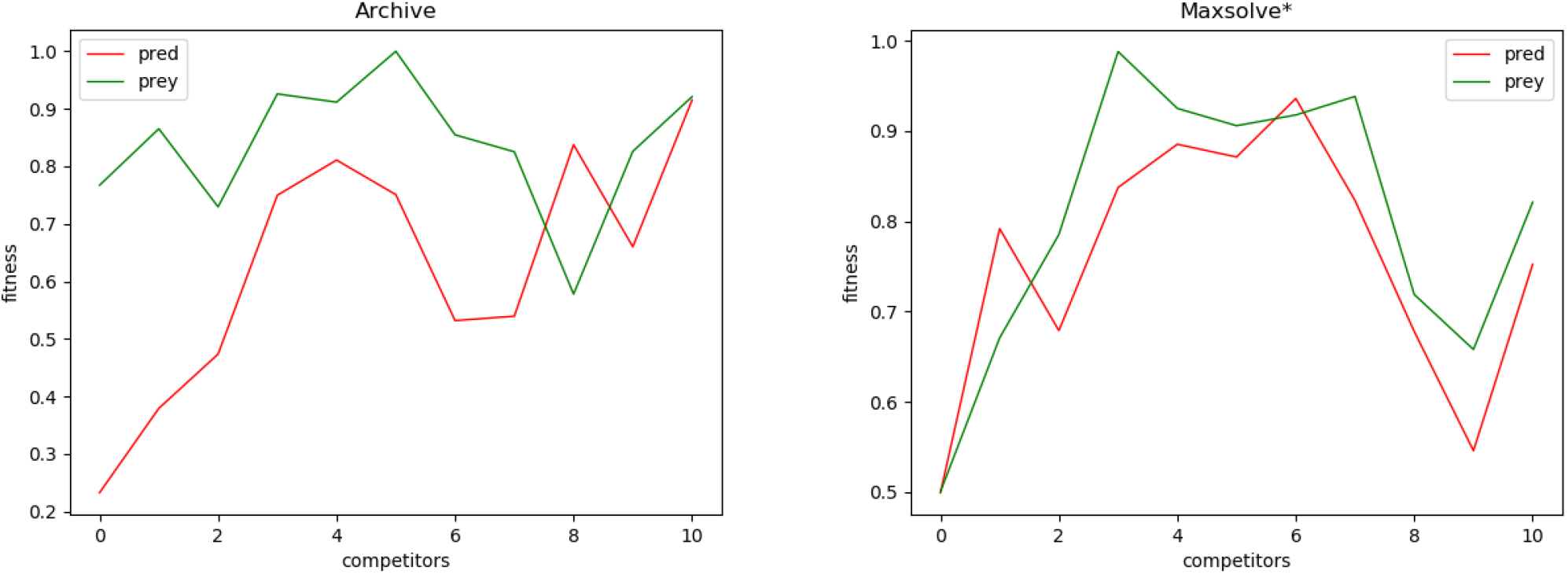
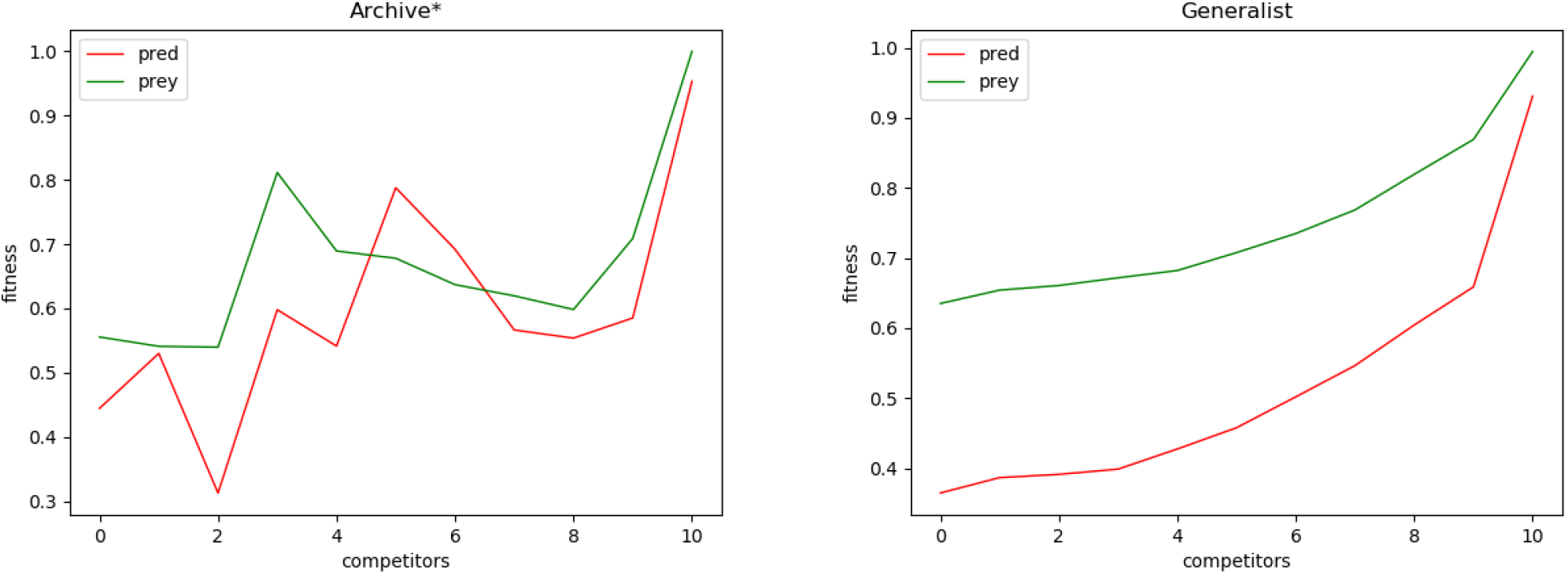
Performance of predators and preys from the last generation against opponents from the same generation (0) and opponents from the 10 preceding stages (1-10), where 10 corresponds to opponents from the initial generation. The 10 stages are separated by 1/10 of the total generations. Results were obtained using the Archive, Maxsolve*, Archive*, and Generalist algorithms. Each plot represents the average results of 10 evolutionary experiments. These plots display the same data as shown in the last row and last columns of the matrices presented in Figure 1.

To identify the best method, we conducted cross-tests comparing the champions obtained from each algorithm against those obtained from other algorithms. For each replication, we selected the best predator and prey among the last 40 evolved individuals in each run. Consequently, we have 10 champion predators and 10 champion preys for each experimental condition. The cross-tests are realized by comparing the performance of a set of 10 champion agents evolved under one experimental condition against champion opponents evolved under the same or a different experimental condition. More specifically, cross-tests values are computed with the following equation:

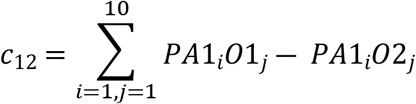

where *c* is the cross-test value, *1* and *2* denote the experimental conditions being compared, *P* indicates the performance, *A* represents the agents, *O* represents the opponents, *i* and *j* are indices for the 10 champion agents and opponents, respectively. Positive and negative cross-test values indicate the superiority and inferiority, respectively, of the first experimental condition over the second.

Table 1 presents the results of cross-tests conducted among the four experimental conditions. Notably, agents evolved using the Archive* method outperform those evolved with the Archive method, for both predators and preys. Furthermore, agents evolved using the Generalist method outperform agents from the other three methods, again for both predators and preys.

**Table 1.** The cross-test of champion agents conducted using the Archive, Maxsolve*, Archive*, and Generalist algorithms. In each row, we evaluate the performance of agents evolved under the condition indicated in that row against opponents evolved under the condition indicated in the corresponding column. We then subtract the performance obtained by evaluating the agents against the opponents indicated in the same row. Positive values indicate that the condition indicated in the row outperforms the condition indicated in the column. The numbers denoted by ‘p=‘ represent the probability that the performance obtained against the two sets of opponents belongs to the same distribution. Values in bold indicate cases where the difference in performance is statistically significant (Mann–Whitney U-test with Bonferroni correction, p-value < 0.0167). The table is divided into two parts: the top section displays cross-tests using predators as agents and preys as opponents, while the bottom section reverses the roles.

| predators |  |  |  |  |
| --- | --- | --- | --- | --- |
|  | Archive | Maxsolve* | Archive* | Generalist |
| Archive |  |  |  |  |
| MaxSolve* | 0.05, p=.270 |  |  |  |
| Archive* | **0.15, p=.001** | 0.04, p=.112 |  |  |
| Generalist | **0.12, p=.007** | **0.13, p=.001** | **0.14, p=.001** |  |

**Table 1.** The cross-test of champion agents conducted using the Archive, Maxsolve\*, Archive\*, and Generalist algorithms.
| preys |  |  |  |  |
| --- | --- | --- | --- | --- |
|  | Archive | Maxsolve* | Archive* | Generalist |
| Archive |  |  |  |  |
| MaxSolve* | 0.04, p=.933 |  |  |  |
| Archive* | **0.17, p=.003** | 0.12, p=.044 |  |  |
| Generalist | **0.27, p=.001** | **0.23, p=.001** | **0.24 p=0.001** |  |

To further validate the effectiveness of the alternative methods and assess the generality of the solutions, we conducted an additional analysis. Specifically, we evaluated the 40 champion predators (obtained from each method in corresponding 10 replications) against the 40 champion preys. The results, displayed in Figure 3, reveal significant differences. Notably, Generalist predator and prey champions outperform the champions obtained with all other methods (Mann–Whitney U-test with Bonferroni correction, p-value < 0.0167). Moreover, Archive* predator and prey champions outperform Archive predators and preys (Mann–Whitney U-test with Bonferroni correction, p-value < 0.0167). Instead, the performances of Maxsolve* and Archive predator and prey champions do not significantly differ (Mann–Whitney U-test with Bonferroni correction, p-value > 0.0167).

**Figure 3.**
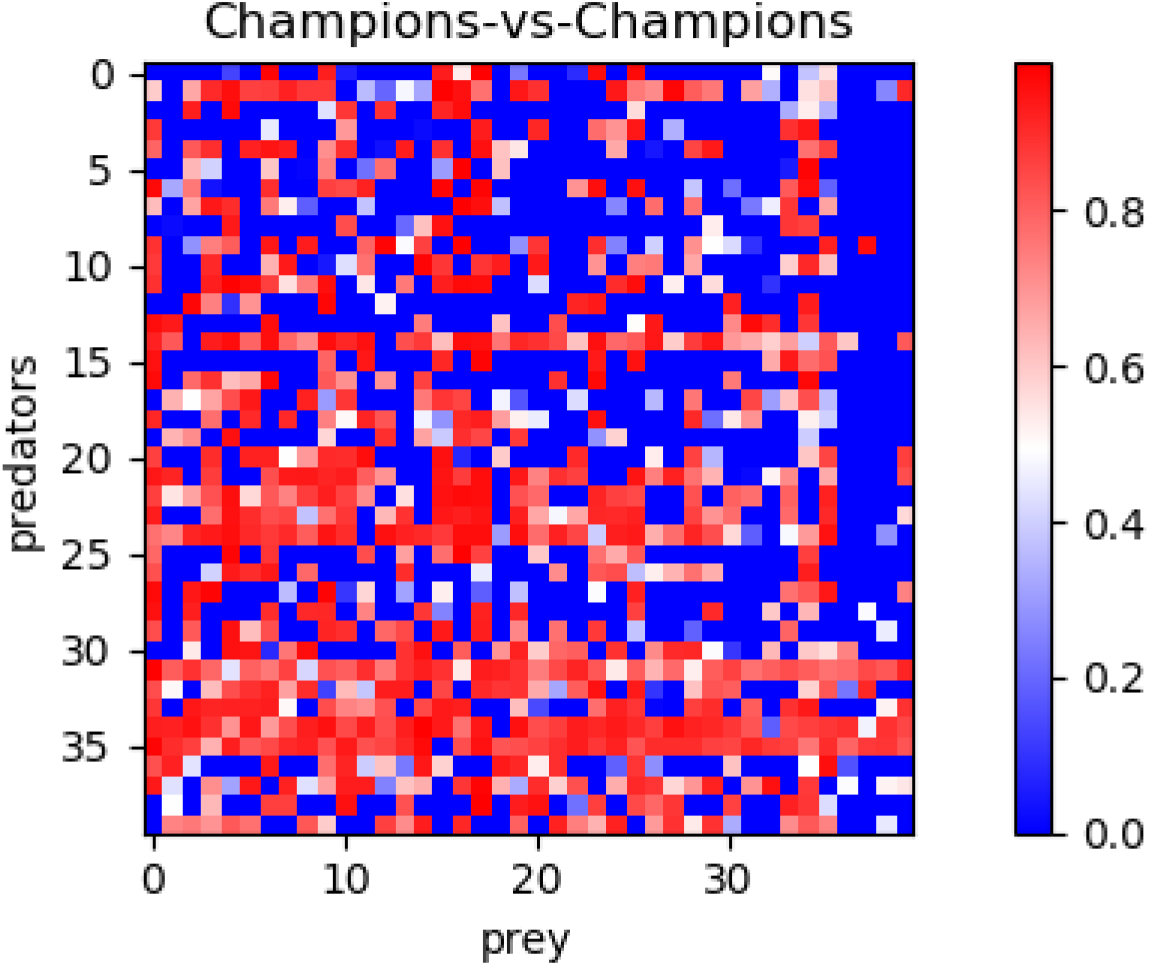
Post-evaluation of the 40 predator champions obtained using four different methods against the 40 prey champions obtained using the same methods. The table consists of four sets of 10 rows and columns, representing the performance of champions from the Archive, Maxsolve*, Archive*, and Generalist methods in their corresponding 10 replications. Each pixel’s color indicates the predator’s performance, while the prey’s performance corresponds to the inverse of the predator’s one.

As shown in Figure 3, the predators and preys obtained using the Generalist method (displayed in rows and columns 30-39, respectively) achieve the best performance. Notably, the fifth and sixth predator champions obtained with the Generalist algorithm (Figure 3, rows 34 and 35) achieve a performance of at least 0.5 against 29 out of the 30 prey champions obtained using the other three methods. Additionally, the seventh and eighth prey champions obtained with the generalist algorithm (Figure 4, columns 36 and 37) also achieve a performance of at least 0.5 against 29 out of 30 predator champions obtained using the other three methods.

The analysis of the behavior displayed by the champions reveals their acquisition of sophisticated behavioral skills. Specifically, some of the champions demonstrate the ability to move both toward the front and rear directions, skillfully alternating their direction of motion based on the circumstances (as shown in Video 1, Appendix C). They exhibit the capability to capture and evade a wide range of opponents. Moreover, they remain robust against adversarial behaviors exhibited by opponents in most cases; in other words, they are rarely fooled by opponent strategies, despite those strategies have been fine-tuned against them. The ability to alternate their direction of motion depending on the circumstances is more commonly observed in agents evolved using the Generalist algorithm.

An illustrative example of an agent vulnerable to specific opponent behavior is the 8th champion predator obtained using the Generalist method. While this predator is generally effective, it proves fragile when confronted with the adversarial strategy employed by the 7th champion prey. This prey displays an oscillatory behavior that triggers a harmless spinning-in-place response from the predator (see Video 2, Appendix C). Another instance of an agent vulnerable to specific opponent behavior is the predator shown in Video 3 (Appendix C). The 4th prey champion, obtained through the Archive* algorithm, effectively neutralizes this specific predator by moving counterclockwise around it, consistently eliciting the same avoidable attacking behavior in the opponent.

## Conclusion

In this article, we delve into the conditions that drive competitive evolution toward genuine progress, i.e. toward solutions that become better and better against all possible opponents. Specifically, we introduced a set of methods for measuring historical and global progress, we discussed factors that facilitate genuine progress, and we compared the efficacy of four algorithms.

The methods considered were the follows: (1) the Archive algorithm (Rosin & Belew, 1997) that promotes global progress by maintaining an archive of the best individuals from previous generations. This permits to evaluate evolving individuals against opponents of current and previous generations. (2) The Maxsolve* algorithm, i.e. a variation of the original De Jong’s algorithm (De Jong, 2005) adapted for both transitive and non-transitive problems. This method also relies on an archive that, however, is used to preserve only diversified individuals. (3) The Archive* algorithm, introduced in this paper, which extends the vanilla Archive method by leveraging multiple evolving populations. This extension allows for the inclusion of more diverse individuals in the archive. (4) The Generalist algorithm (Simione & Nolfi, 2021) that does not use an archive but incorporates a mechanism for identifying and discarding variations leading to local progress only.

The results obtained in a predator-prey scenario, commonly used to study competitive co-evolution, demonstrate that all the considered methods lead to global progress in the long term. However, the rate of progress and the ratio of progress versus retrogressions vary significantly.

The Generalist method outperforms the other three methods and is the only one capable of producing solutions that consistently score better and better across generations against previous and future opponents in successive evolutionary phases. The other three methods also exhibit retrogression phases, although less frequently than progression phases. Additionally, the Archive* algorithm, introduced in this paper, outperforms both the vanilla Archive Algorithm and the MaxSolve* algorithm.

The superiority of the Generalist algorithm is also demonstrated through visual comparisons of the behavior exhibited by the evolving robots. Indeed, the ability to move bi-directionally and appropriately alternate the direction of motion depending on the circumstances, providing a significant advantage, is more commonly observed among the robots evolved with the Generalist algorithm than among the robots evolved with other algorithms.

Future research should verify whether our results generalize to other competitive settings and whether the continuation of evolutionary progress can lead to an open-ended dynamic in which the efficacy of the evolved solutions keeps increasing in an unbounded manner.

## Appendix A

Below we include the pseudo-code of the algorithms. In all cases the connection weights of the controllers of the robots are evolved by using the OpenAI-ES algorithm (Salimans et al., 2017) and by using the hyperparameters indicated in the reference above. More specifically, observation vectors were normalized by using virtual batch normalization (Salimans et al, 2016, 2017), the connections weights were normalized by using weight decay, the distribution of the perturbations of parameters was set to 0.02, and the step size of the Adam optimizer was set to 0.01. The fitness gradient is estimated by generating 40 offspring, i.e. 40 perturbed versions of the parent. The fitness of the evolving individuals corresponds to the average fitness obtained during 10 episodes in which they are evaluated against 10 different opponents in 10 corresponding evaluation episodes. The fitness and the perturbation vectors are used to compute the gradient of the expected fitness, which is used to update the parameters of the parent through the Adam (Kingma, & Ba, 2014) stochastic optimizer. The predator and prey robots are evolved in parallel. The evolutionary process is continued until the total number of evaluation steps performed exceeds 75 • 10^9^.

In the case of the Archive and Maxsolve* algorithms, the parameters of the evolving robots are generated from a single parent (a parent for the predator and a parent for the prey robots). In the case of the Archive* algorithm, they are generated from 10 parents. The generalist algorithm, instead, uses a population of 80 parents, evolves 10 candidate parents generated by creating a copy of 10 parents selected randomly every 20 generations, and replace the worst parents with the candidate parents outperforming them. The performance of parents and of candidate parents are computed by evaluating the parents against the full set of 80 opponents, i.e. by evaluating the candidate parents also against opponents not encountered during their evolution. The steps performed to evaluate performance contribute to increase the total number of steps performed.

The Archive and Archive* algorithms preserve the best robots of each generation in archives that keep increasing in size during the evolutionary process. The Maxsolve* algorithm uses archives that can grow up to a maximum size only. This is realized by: (i) storing in a fitness_data[X][Y] matrix the average fitness obtained by agents of generation X against opponents of generation Y, (ii) computing the average fitness obtained by agents contained in the archive against opponents of 10 subsequent evolutionary phases, and (iii) eliminating one dominated agent selected randomly from the archive in each generation. An agent is dominated when the average fitness obtained against opponents of different phases is consistently equal or lower than the average fitness obtained by another agent included in the archive.

### Archive algorithm

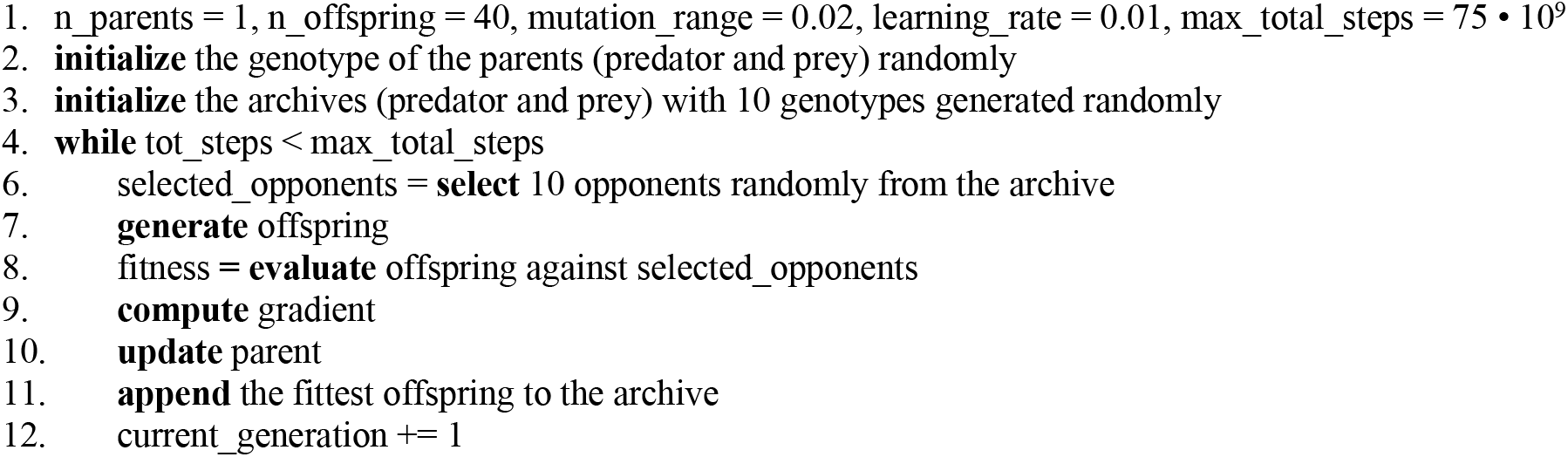

### Maxsolve* algorithm

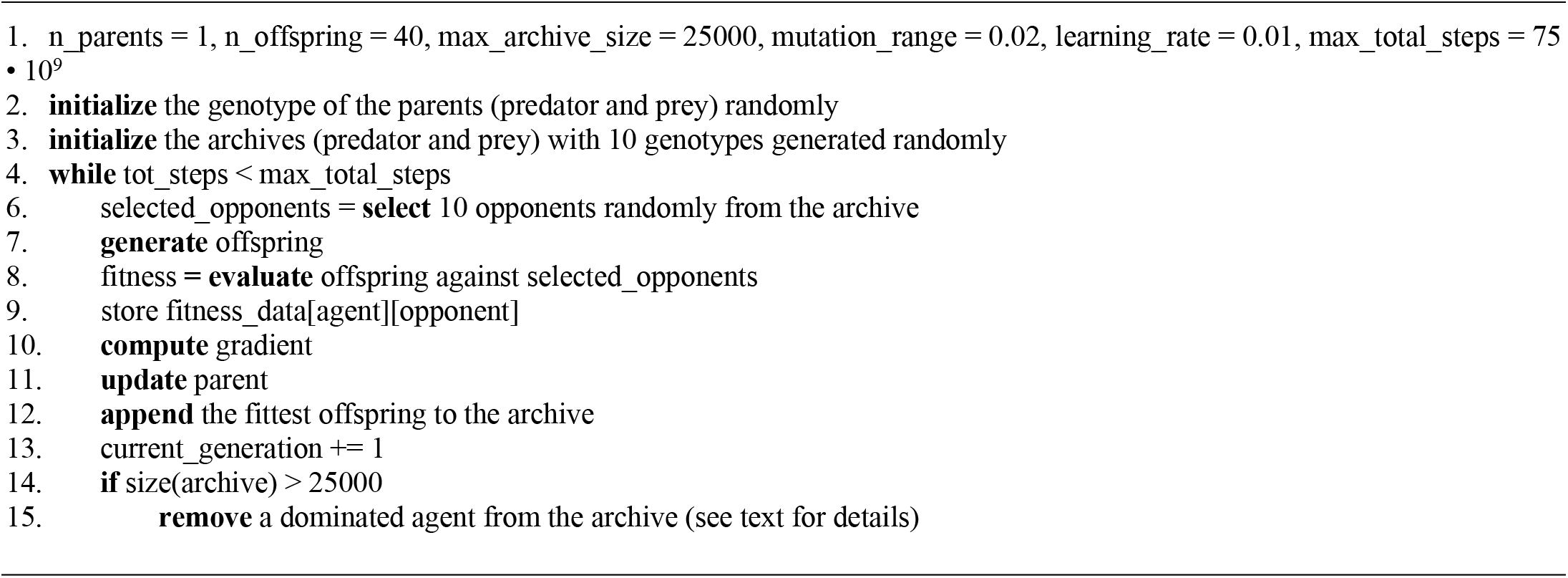

### Archive* algorithm

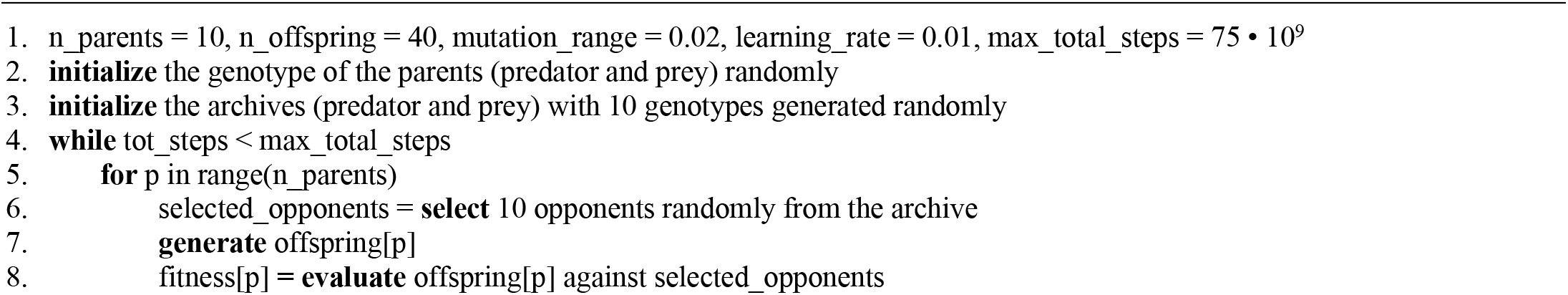
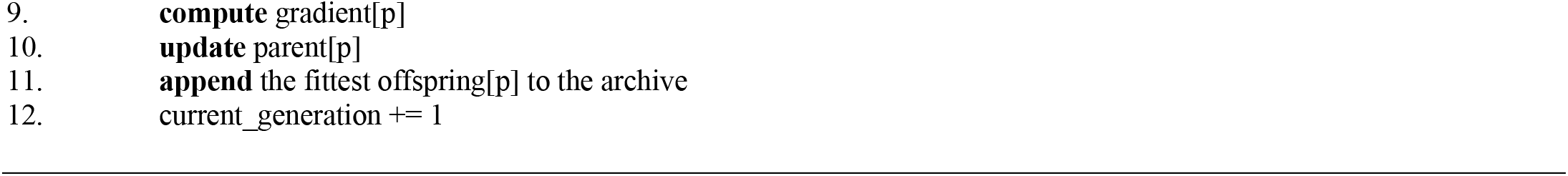

### Generalist algorithm

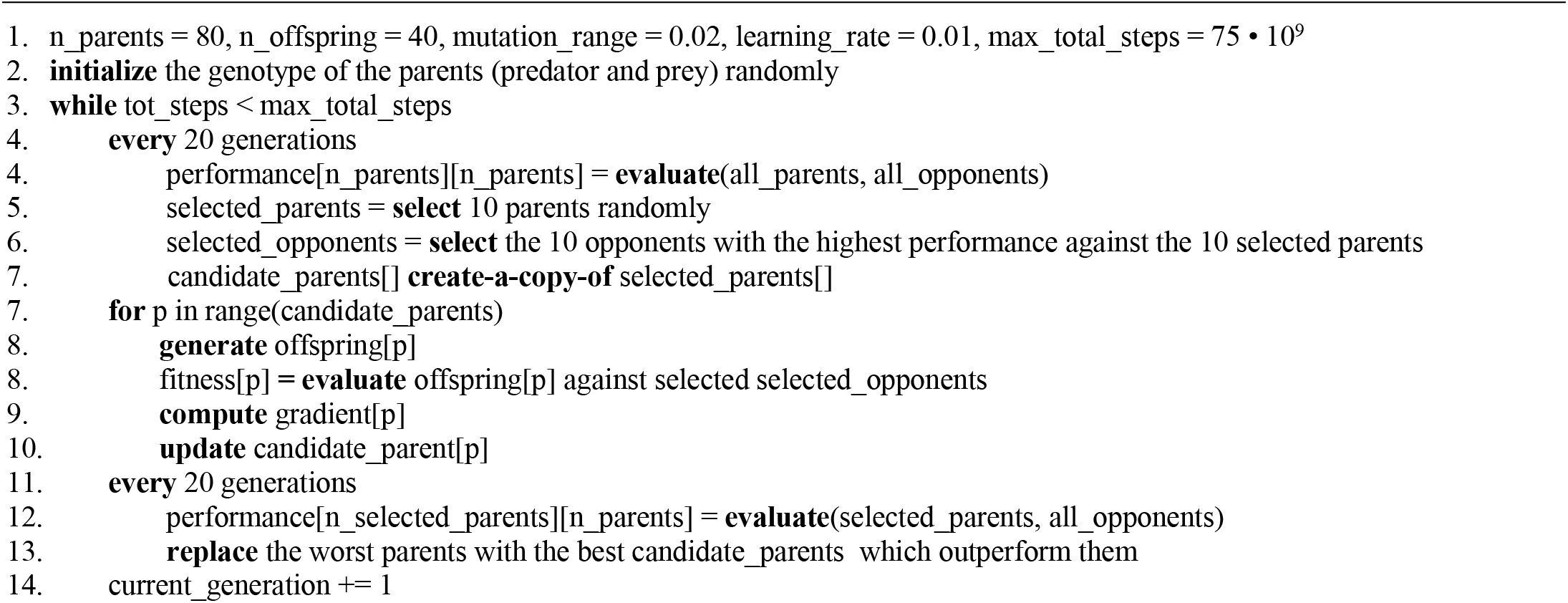

## Appendix B

The robots were simulated MarXbots (Bonani et al., 2010), i.e. circular robots with a 17cm diameter. They are equipped with a differential drive motion system, a ring of 24 color LEDs, 24 infrared sensors, 4 ground sensors, an omnidirectional camera, and a traction sensor. In the experiments, the LEDs of predator robots are set in red, while the LEDs of prey robots are set in green. The robots were placed in a 3×3 m square arena surrounded by black walls. The arena floor was grayscale, varying from white to black from the center to the periphery (see Fig. A1).

The maximum wheel speed that the wheels of the differential drive motion system could assume was 10 and 8.5 rad/s, for the preys and the predators respectively. The relative speed of the two robot types was adjusted to balance the overall complexity of the problem faced by predators and preys, i.e. preventing one species from reaching maximum or minimum fitness. The environment state, robot sensors, motors, and neural network were updated at a frequency of 10 Hz.

The neural network controller of the robot consists of a LSTM (Long Short-Term Memory, see Hochreiter & Schmidhuber, 1997; Gers & Schmidhuber, 2001) recurrent neural network with 23 sensory neurons, 25 internal units, and 2 motor neurons. The sensory layer includes 8 sensory neurons encoding the average activation state of eight groups of three adjacent infrared sensors, 8 neurons that encode the fraction of green or red light perceived in the eight 45° sectors of the visual field of the camera, 1 neuron that encodes the average amount of green or red light detected in the entire visual field of the camera, 4 neurons encoding the state of the four ground sensors, 1 neuron that encodes the average activation of the four ground sensors, 1 neuron that encodes whether the robot collides with an obstacle (i.e., whether the traction force detected by the traction sensor exceeds a threshold). The sensory neuron states are normalized in the range [0.0, 1.0]. The motor layer includes two neurons encoding the desired translational and rotational motion of the robot in the range [0.0, 1.0].

**Fig. A1.**
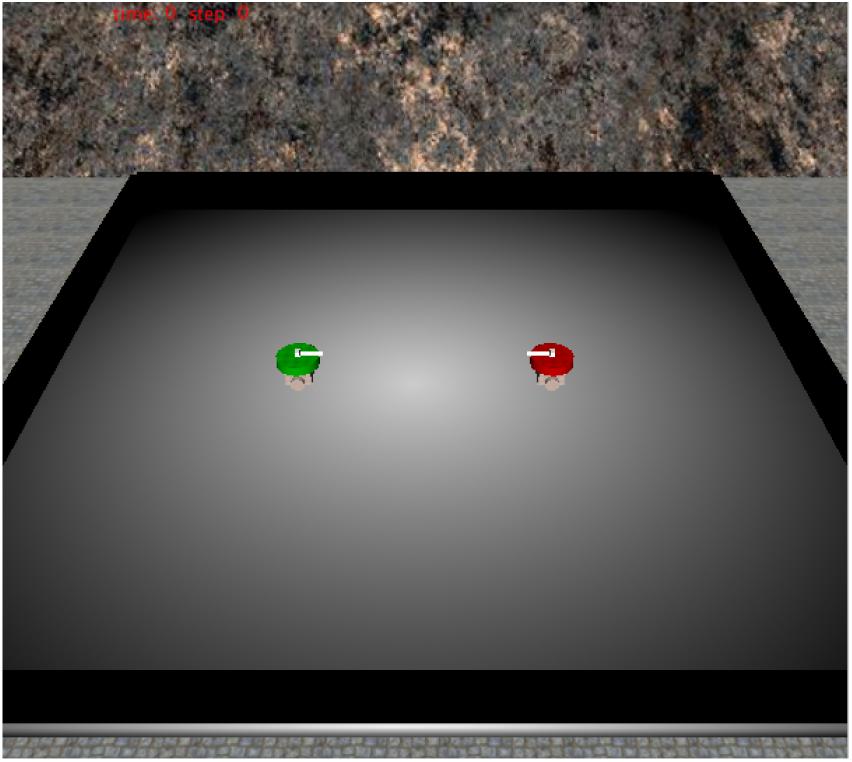
The robots and the environment in simulation. The red and green robots correspond to the predator and prey robots, respectively.

## Appendix C

**Video 1**. Available from https://youtube.com/shorts/prpFwtN-v3Y?feature=share

**Video 2**. Available from https://youtube.com/shorts/77mYMT6CKnI?feature=share

**Video 3**. Available from https://youtube.com/shorts/hvrs2rcJVdU?feature=share

## Acknowledgment

I acknowledge financial support from PNRR MUR project PE0000013-FAIR.

